# A parsimonious software sensor for estimating the individual dynamic pattern of methane emissions from cattle

**DOI:** 10.1101/298679

**Authors:** R. Muñoz-Tamayo, J. F. Ramírez Agudelo, R. J. Dewhurst, G. Miller, T. Vernon, H. Kettle

**Affiliations:** UMR Modélisation Systémique Appliquée aux Ruminants, INRA, AgroParisTech, Université Paris-Saclay, 75005, Paris, France; Universidad de Antioquia - UdeA, Facultad de Ciencias Agrarias. Grupo de Investigación en Ciencias Agrarias -GRICA. Ciudadela de Robledo, Carrera 75 Nº 65·87, Medellín, Colombia; Future Farming Systems, SRUC, West Mains Road, Edinburgh, EH9 3JG, United Kingdom; Biomathematics and Statistics Scotland (BioSS), Kings Buildings, Edinburgh, EH9 3FD, United Kingdom

**Author notes:** Corresponding author: Rafael Muñoz-Tamayo.

**Keywords:** greenhouse gas, methane, modelling, ruminant, precision farming

## Abstract

Large efforts have been deployed in developing methods to estimate methane emissions from cattle. For large scale applications, accurate and inexpensive methane predictors are required. Within a livestock precision farming context, the objective of this work was to integrate real-time data on animal feeding behaviour with an *in silico* model for predicting the individual dynamic pattern of methane emission in cattle. The integration of real-time data with a mathematical model to predict variables that are not directly measured constitutes a software sensor. We developed a dynamic parsimonious grey-box model that uses as predictor variables either dry matter intake (DMI) or the intake time (IT). The model is described by ordinary differential equations. Model building was supported by experimental data of methane emissions from respiration chambers. The data set comes from a study with finishing beef steers (cross-bred Charolais and purebred Luing finishing). DMI and IT were recorded with load cells. A total of 37 individual dynamic patterns of methane production were analysed. Model performance was assessed by concordance analysis between the predicted methane output and the methane measured in respiration chambers. The model predictors DMI and IT performed similarly with a Lin’s concordance correlation coefficient (CCC) of 0.78 on average. When predicting the daily methane production, the CCC was 0.99 for both DMI and IT predictors. Consequently, on the basis of concordance analysis, our model performs very well compared with reported literature results for methane proxies and predictive models. Since IT measurements are easier to obtain than DMI measurements, this study suggests that a software sensor that integrates our *in silico* model with a real-time sensor providing accurate IT measurements is a viable solution for predicting methane output in a large scale context.

**Implications:** Reducing methane emissions from ruminants is a major target for sustainable and efficient livestock farming. For the animal, methane production represents a loss of feed energy. For the environment, methane exerts a potent greenhouse effect. Methane mitigation strategies require accurate, non-invasive and inexpensive techniques for estimating individual methane emissions on farm. In this study, we integrate measurements of feeding behaviour in cattle and a mathematical model to estimate individual methane production. Together, model and measurements form a software sensor that efficiently predicts methane output. Our software sensor is a promising approach for estimating methane emissions at large scale.

## Introduction

Methane emission from cattle is an output associated to animal efficiency that impacts the environmental footprint of livestock farming. Accordingly, reducing enteric methane production is a major target for ruminant production systems (Martin *et al.*, 2010, Hristov *et al.*, 2013). Large efforts have been deployed to develop methods and devices to measure and estimate methane emissions from ruminants, with respiration chambers being the gold standard under rigorous operation (Gardiner *et al.*, 2015, Hammond *et al.*, 2016). However, some of these techniques are usually costly and not suitable to be applied for on farm application at large scale for the development of mitigation strategies of greenhouse gas emissions. An ideal technique for large scale application should provide, at low cost, individual accurate estimations of methane produced by ruminants (Negussie *et al.*, 2017b). In complement to the development of methane proxies, mathematical modelling offers a useful tool for methane prediction. Mathematical models are often categorized as white box (phenomenological, mechanistic) or black box (empirical) models. A model with mechanistic and empirical components is termed as a grey box model. With respect to the models developed for predicting methane production by ruminants, white box models aim at describing the biological phenomena associated to rumen fermentation and methanogenesis (*Mills et al., 2001*, Huhtanen *et al.*, 2015, Vetharaniam *et al.*, 2015). These phenomena may include for instance the microbial activity of archaea methanogens (Wang *et al.*, 2015, Muñoz-Tamayo *et al.*, 2016). Alternatively, black box models aim at deriving regression equations that quantify relationships between variable predictors and methane emissions (Sauvant *et al.*, 2011, Ramin and Huhtanen, 2013). In general, white box models offer the possibility of quantifying the dynamics of key variables while black box models are often static. On the other hand, black box models are less complex than white box model which favours their implementation for practical purposes (*e.g.*, on-farm monitoring). Existing black box models for methane predictions are algebraic equations that use an average measure of dry matter intake (DMI) as primary predictor (Giger-Reverdin *et al.*, 2003, Charmley *et al.*, 2016, Niu *et al.*, 2018). Generally, models and techniques have been applied to estimate the daily average methane emission. Few studies report predictions of the dynamic pattern of methane production (*Wang et al., 2015*). Integrating dynamic data from dedicated sensors with mathematical models to support livestock management decisions, and guide timely interventions is the great promise of precision livestock farming (Wathes *et al.*, 2008, Rutten *et al.*, 2013, Friggens *et al.*, 2017). The integration of real-time data with a mathematical model to predict variables that are not directly measured constitutes what is called as software sensor (observer) in the automatic control scientific literature (Dochain, 2003). Software sensors have been broadly applied to monitor and control biotechnological processes. A highly performant software sensor is composed of (i) real-time sensors that accurately measure variables of interest and (ii) a reliable model that provides accurate predictions and has a simple structure to facilitate its implementation. In this context, the objective of this work was to develop a software sensor for predicting the individual dynamic pattern of methane emissions in cattle. While, a software sensor operates in real-time using online sensor measurements, for a research purpose, our development was applied to off-line data as preliminary step for further on-line applications. Our software sensor is composed of a dynamic grey box model and dynamic data on animal feeding behaviour measured either as dry matter intake (DMI) or simply as intake time (IT). Our developments provide a promising and viable solution for predicting methane output for on farm applications at large scale.

## Material and methods

### Experimental data

Model building was supported by the analysis of experimental data obtained from studies conducted at Scotland’s Rural College (SRUC, United Kingdom) with finishing beef steers from two breeds (cross-bred Charolais and purebred Luing) (Troy *et al.*, 2015). Animals received two contrasting basal diets consisting (g/kg DM) of 500:500 and 80:920 forage to concentrate ratios. Within each basal diet, there were two treatments: a control treatment with rapeseed meal as protein source, and an oil treatment with rapeseed cake as protein source to increase dietary oil from 27 (control) to 53 g/kg DM. Methane emissions were measured in a respiration chamber facility with a turnover rate constant of 0.04 min^-1^ and a gas recovery of 98% (Rooke *et al.*, 2014). The gas sampling time was 6 min. DMI and IT were recorded with load cells. A total of 37 individual dynamic patterns of methane production was analysed.

### Mathematical model development

A mass balance applied to the respiration chamber for methane gives the following ordinary differential equation (ODE)

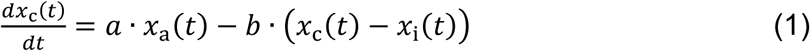

Where *x*_c_, *x*_i_ are the amount (in grams) of methane inside the chamber and at the inlet of the chamber respectively, and *x*_a_ is the amount of methane in the gas flow released by the animal (exhalation+ eructation). The parameter *a* (min^-1^) is the rate constant of the animal gas emission and *b* (min^-1^) is the turnover rate of the chamber. Note that in reality, *a* may be time varying. If *x*_i_ is almost constant over time and *x*_i_ ≪ *x*_c_, then Eq. (1) is simplified to

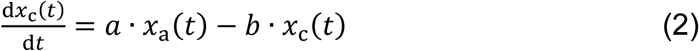

The quantity *a* ∙ *x*_a_ is the gas produced (g/min) by the animal while *b* ∙ *x*_c_ is the gas production (g/min) measured in the chamber. For mathematical convenience, we denote *y*_a_ = *a* ∙ *x*_a_ and *y*_c_ = *b* ∙ *x*_c_. Equation (2) is thus translated to

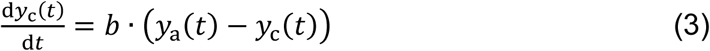

If the turnover rate of the chamber is optimally chosen, it can be shown that *y*_c_ follows almost the same dynamics of *Y*_a_ (See Supplementary material S1). In the remaining of the text, we will assume that *Y*_c_ = *Y*_a_. Based on this clarification, we proposed the following ODE model for predicting the animal methane emission *Y*_a_

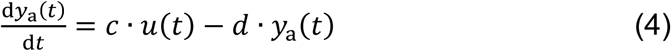

where *u* is either the DMI or IT. DMI is in g/min and IT is a Boolean variable having the value one to indicate intake (eating) activity and zero otherwise. The parameters *c,d* are specific to the animal and diet and must be estimated from the experimental data. The parameter *d* is in min^-1^. The parameter *c* is in g CH_4_/(g DM∙min) or in g CH_4_/min^2^ when using DMI or IT as predictors respectively. The model in Eq. (4) has a parsimonious structure with only two parameters. Although very simple, it follows the structure of a mass balance model (as Eq. (2)) in aggregated form. Indeed, the quantity parameter / can be interpreted as a yield factor *i.e*, the mass of methane produced per mass of DM (when using DMI as a predictor). Given this phenomenological characteristic, the model is referred to as a grey box model. Additionally, the model has the property of being identifiable, that is that the parameters *c,d* can, in theory, be uniquely estimated if noise-free dynamic data of *Y*_a_ and *u* are available (see, *e.g.*, Muñoz-Tamayo *et al.*, 2018 for a discussion on parameter identifiability). The model in Eq. (4) can also be written in finite differential form. By applying backward differentiation with a constant time step ∆*t*, we obtain

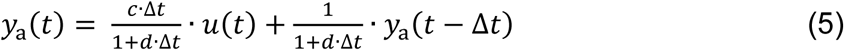

Equation (5) is an exponential smoothing filter. After a sensitivity analysis (not shown) the step time was fixed to ∆*t* = 1.0 min. The model was implemented in the open source software Scilab (https://www.scilab.org). Model calibration was performed by minimizing the sum of squared errors beween experimental data and predicted output for each of the 37 dynamic methane patterns. The minimization was performed using the Nelder-Mead algorithm implemented in the fminsearch function of Scilab. Our grey box model has the simplest structure to represent the dynamics of methane emissions from time series data of DMI or IT. To assess if increasing model complexity could lead to gains in goodness of fitting, we tested the performance of different linear models (described by Laplace transfer functions) with higher number of parameters than our model using the Matlab® System Identification Toolbox (Ljung, 1997). Our model was the best linear candidate model with respect to the Akaike’s information criterion which provides an indicator of model parsimony based on a trade-off between goodness of fit and model complexity (quantified by the number of model parameters). The Lin’s concordance correlation coefficient (Lin, 1989) was computed to quantify the agreement between the methane estimation provided by the software sensor and the methane measured in respiration chambers (the gold standard).

## Results

Figure 1 shows typical data extracted from the experimental study. The dynamics of methane production is modulated by the feeding pattern (DMI or IT). Methane emissions increased following feeding and declines towards a basal value before the next feeding as observed in other studies (Crompton *et al.*, 2011, Wang *et al.*, 2015, Olijhoek *et al.*, 2016).

**Figure 1.**
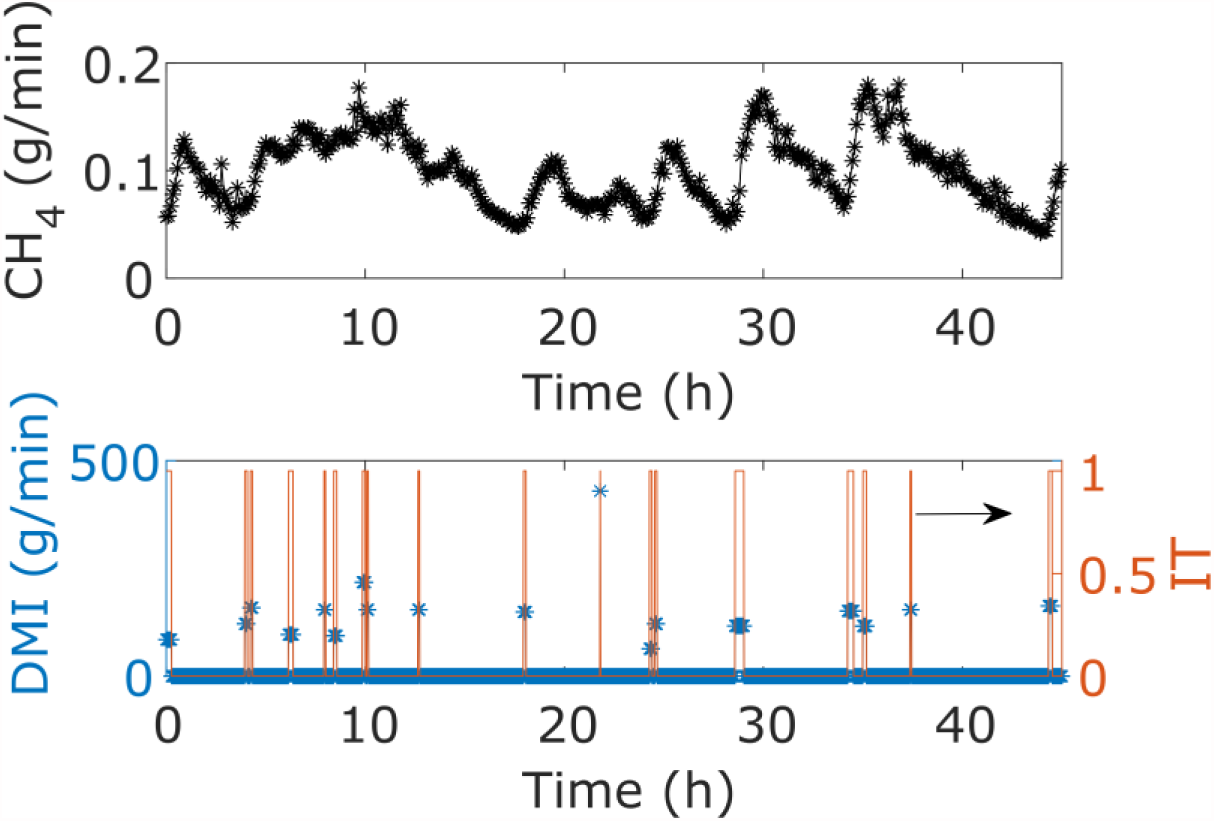
Example of dynamic data of methane production (top) and feeding behaviour measured as DMI (*) and IT (solid line) Figure 2 displays the individual dynamic pattern of methane production against software sensor predictions for the best and worst fitting cases. Plots are given for the model using either DMI or IT as predictors applied to both control and oil treatments.

**Figure 2.**
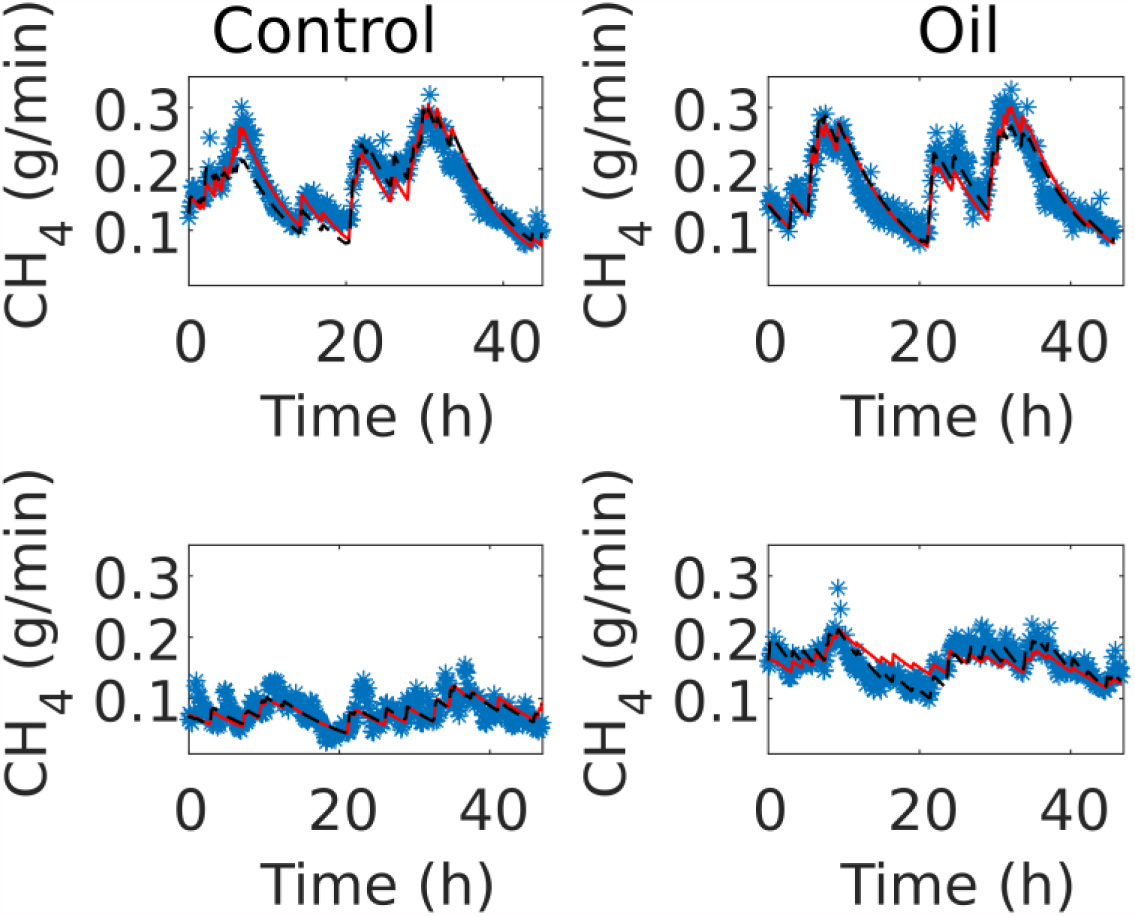
Experimental (*) versus predicted methane emissions using DMI (red solid line) and IT (dashed black line) as predictors for control and oil treatments. Top plots are the experiments where model fits were the best. Bottom plots are the experiments where model fits were the poorest. IT is as good predictor as DMI

**Figure 3.**
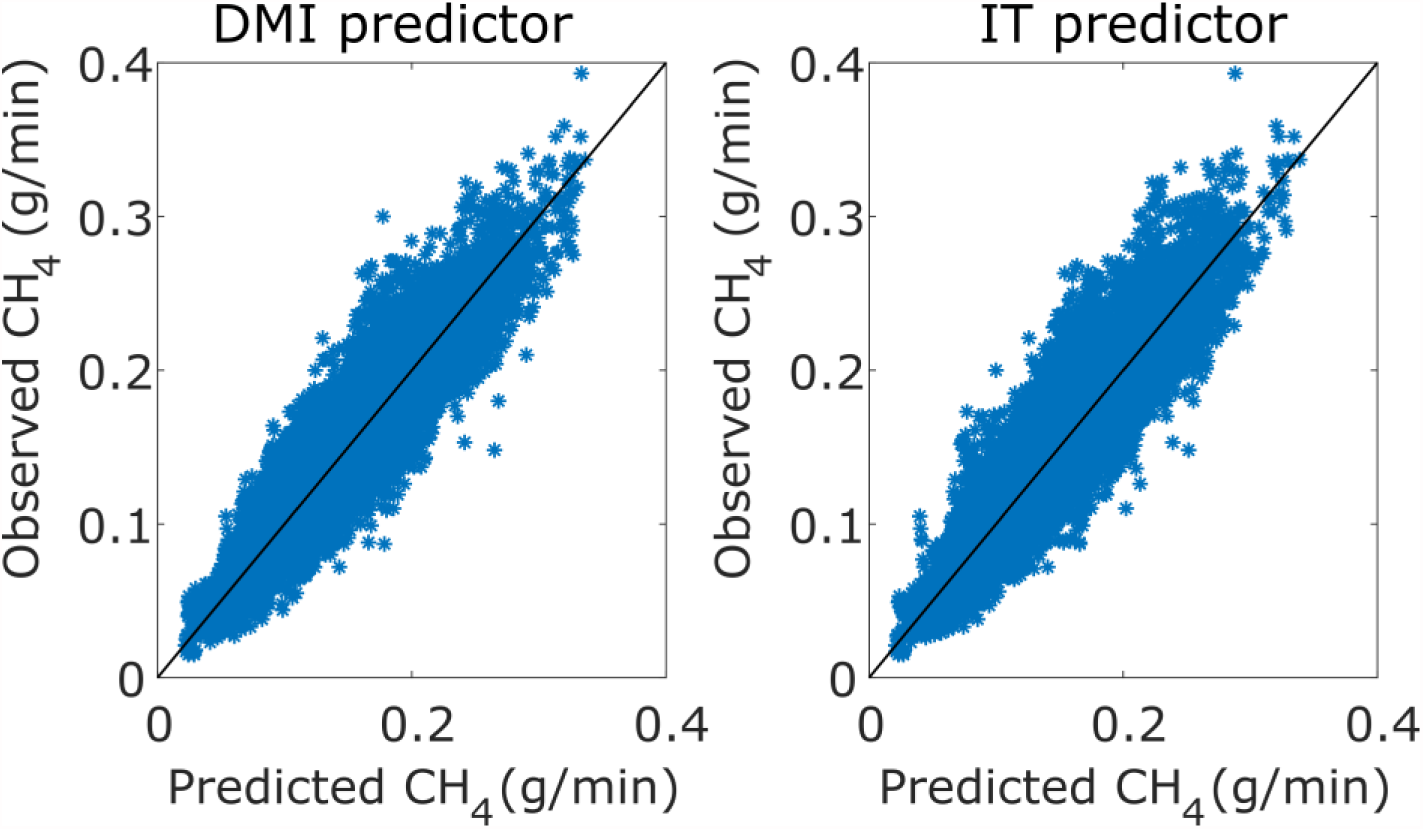
Experimental data vs predicted output of the dynamic pattern of methane of production. The isocline is the solid line. Results are presented for the model using either DMI or IT as predictors Figure 3 displays the observations versus predictions from both models for all dynamic individual data (n = 15041 time data points). Figure 4 shows the individual daily average methane emission (n=37 steers) against predicted methane production. It is observed that individuals fed with the mixed basal diet produce more methane than those fed with the control basal diet (Troy *et al.*, 2015).

**Figure 4.**
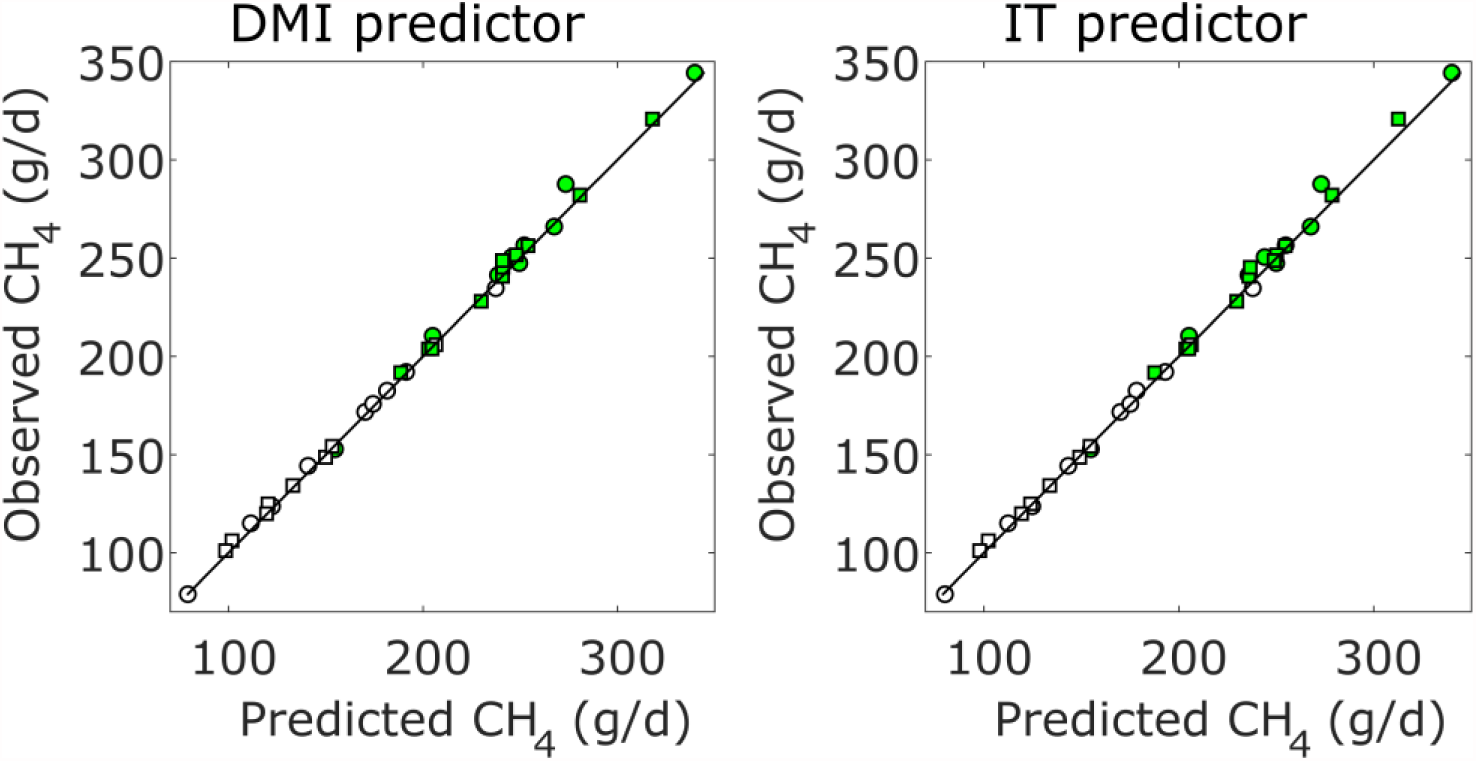
Experimental data vs predicted output of daily average methane emission production for control (o) and oil (□) treatments. Filled marks are for the mixed basal diet, unfilled marks are for the concentrate basal diet. The isocline is the solid line. Results are presented for the model using either DMI or IT as predictors

Tables 1 and 2 show the results of the model calibration for the individual dynamics of methane production using either DMI or IT as predictors for the control and oil treatments respectively. Classical statistical indicators are also given. The statistical analysis about the effects of genotype, basal diet and treatment on methane production has already been published (Troy *et al.*, 2015). To avoid redundancy, we focus here on the analysis of model parameters.

**Table 1.** Model calibration results of individuals for the control treatment

| Animal* | Model with DMI as predictor |  |  |  |  | Model with IT as predictor |  |  |  |  |
| --- | --- | --- | --- | --- | --- | --- | --- | --- | --- | --- |
| | $c \cdot 10^{-5}$ | $d \cdot 10^{-3}$ | CCC** | $r^2$ | CV<sub>RMSE</sub>*** | $c \cdot 10^{-3}$ | $d \cdot 10^{-3}$ | CCC | $r^2$ | CV<sub>RMSE</sub> |
|  | gCH<sub>4</sub>/(g DM·min) | min<sup>-1</sup> |  |  |  | gCH<sub>4</sub>/min<sup>2</sup> | min<sup>-1</sup> |  |  |  |
| ChC1 | 2.93 | 2.35 | 0.74 | 0.51 | 22.53 | 3.47 | 2.12 | 0.78 | 0.61 | 20.07 |
| ChC2 | 2.51 | 2.53 | 0.78 | 0.58 | 23.29 | 2.04 | 2.32 | 0.72 | 0.50 | 25.50 |
| ChC4 | 3.63 | 2.31 | 0.80 | 0.64 | 18.71 | 3.19 | 2.33 | 0.72 | 0.54 | 21.27 |
| ChC5 | 3.55 | 1.38 | 0.85 | 0.72 | 13.81 | 2.36 | 1.49 | 0.80 | 0.61 | 16.38 |
| ChC6 | 2.84 | 1.90 | 0.66 | 0.46 | 14.41 | 2.83 | 1.68 | 0.61 | 0.42 | 14.89 |
| ChM1 | 4.80 | 2.12 | 0.92 | 0.83 | 12.90 | 3.78 | 1.90 | 0.89 | 0.77 | 14.89 |
| ChM3 | 3.08 | 1.17 | 0.82 | 0.63 | 16.55 | 0.79 | 0.47 | 0.70 | 0.50 | 19.21 |
| ChM4 | 4.53 | 1.49 | 0.81 | 0.63 | 12.03 | 7.22 | 1.87 | 0.85 | 0.69 | 10.96 |
| ChM5 | 4.17 | 1.62 | 0.77 | 0.59 | 16.70 | 3.60 | 1.46 | 0.77 | 0.57 | 17.07 |
| LuC1 | 3.21 | 2.26 | 0.88 | 0.76 | 13.47 | 2.51 | 1.78 | 0.81 | 0.69 | 15.38 |
| LuC3 | 3.50 | 1.76 | 0.83 | 0.72 | 13.26 | 2.51 | 1.57 | 0.71 | 0.52 | 17.35 |
| LuC4 | 3.41 | 1.60 | 0.83 | 0.67 | 10.61 | 1.88 | 0.98 | 0.77 | 0.63 | 11.26 |
| LuC6 | 2.00 | 1.58 | 0.52 | 0.26 | 24.35 | 2.06 | 1.58 | 0.53 | 0.31 | 23.64 |
| LuM1 | 4.59 | 1.87 | 0.91 | 0.79 | 13.92 | 3.93 | 1.88 | 0.90 | 0.78 | 14.37 |
| LuM3 | 2.21 | 1.24 | 0.61 | 0.43 | 20.09 | 1.74 | 1.08 | 0.59 | 0.41 | 20.35 |
| LuM4 | 4.37 | 1.37 | 0.81 | 0.65 | 16.02 | 5.29 | 1.85 | 0.74 | 0.52 | 18.90 |
| LuM5 | 5.30 | 1.90 | 0.87 | 0.71 | 13.54 | 3.42 | 1.91 | 0.91 | 0.82 | 10.71 |
| LuM6 | 3.52 | 1.41 | 0.89 | 0.79 | 12.70 | 2.48 | 1.46 | 0.87 | 0.74 | 13.93 |
| Mean | 3.56 | 1.77 | 0.79 | 0.63 | 16.05 | 3.06 | 1.65 | 0.76 | 0.59 | 17.01 |

\* Animals are identified by the type of breed: Charolais (Ch) or Luing (Lu), and the type of basal diet: mixed (M) or concentrate (C).
\*\* CCC: Lin's concordance correlation coefficient.
\*\*\* CV<sub>RMSE</sub>: coefficient of variation of the root mean squared error.

**Table 2.** Model calibration results of individuals for the oil treatment

| Animal* | Model with DMI as predictor |  |  |  |  | Model with IT as predictor |  |  |  |  |
| --- | --- | --- | --- | --- | --- | --- | --- | --- | --- | --- |
| | $c \cdot 10^{-5}$ | $d \cdot 10^{-3}$ | CCC** | $r^2$ | CV<sub>RMSE</sub>*** | $c \cdot 10^{-3}$ | $d \cdot 10^{-3}$ | CCC | $r^2$ | CV<sub>RMSE</sub> |
|  | gCH<sub>4</sub>/(g DM·min) | min<sup>-1</sup> |  |  |  | gCH<sub>4</sub>/min<sup>2</sup> | min<sup>-1</sup> |  |  |  |
| ChC1 | 2.90 | 2.08 | 0.88 | 0.77 | 21.01 | 4.20 | 2.36 | 0.89 | 0.79 | 20.33 |
| ChC3 | 2.52 | 1.13 | 0.84 | 0.72 | 12.71 | 1.54 | 1.15 | 0.85 | 0.73 | 12.34 |
| ChC5 | 3.21 | 2.65 | 0.80 | 0.63 | 18.06 | 2.20 | 2.61 | 0.84 | 0.72 | 15.66 |
| ChC6 | 2.94 | 1.54 | 0.76 | 0.52 | 15.94 | 2.12 | 1.49 | 0.87 | 0.76 | 11.20 |
| ChM1 | 2.14 | 0.95 | 0.53 | 0.31 | 14.05 | 2.49 | 1.54 | 0.82 | 0.66 | 9.82 |
| ChM2 | 3.48 | 1.45 | 0.80 | 0.61 | 11.38 | 2.29 | 1.39 | 0.81 | 0.64 | 10.99 |
| ChM3 | 3.28 | 1.39 | 0.82 | 0.66 | 12.45 | 2.88 | 1.10 | 0.69 | 0.46 | 15.73 |
| ChM4 | 3.79 | 1.54 | 0.84 | 0.70 | 13.20 | 3.59 | 1.33 | 0.73 | 0.51 | 16.97 |
| ChM5 | 4.49 | 2.06 | 0.91 | 0.80 | 11.70 | 3.53 | 1.95 | 0.76 | 0.47 | 19.05 |
| ChM6 | 2.77 | 1.40 | 0.86 | 0.77 | 12.75 | 1.85 | 1.19 | 0.83 | 0.73 | 13.89 |
| LuC2 | 1.74 | 1.25 | 0.53 | 0.26 | 14.67 | 0.19 | 0.33 | 0.15 | 0.05 | 16.58 |
| LuC3 | 3.17 | 2.40 | 0.78 | 0.60 | 23.11 | 4.21 | 2.27 | 0.77 | 0.59 | 23.26 |
| LuC4 | 2.73 | 1.77 | 0.87 | 0.76 | 18.44 | 2.27 | 1.79 | 0.89 | 0.80 | 16.62 |
| LuC5 | 3.29 | 2.63 | 0.71 | 0.53 | 18.72 | 2.24 | 2.39 | 0.69 | 0.48 | 19.57 |
| LuM1 | 4.58 | 2.01 | 0.90 | 0.81 | 10.49 | 4.86 | 1.97 | 0.86 | 0.74 | 12.32 |
| LuM2 | 4.74 | 1.95 | 0.91 | 0.82 | 14.59 | 3.57 | 1.88 | 0.93 | 0.86 | 12.93 |
| LuM4 | 4.88 | 2.55 | 0.65 | 0.35 | 20.04 | 7.34 | 2.43 | 0.62 | 0.31 | 20.69 |
| LuM5 | 3.56 | 1.45 | 0.79 | 0.47 | 11.48 | 2.47 | 1.22 | 0.90 | 0.82 | 6.69 |
| LuM6 | 4.34 | 2.17 | 0.75 | 0.61 | 11.97 | 2.36 | 1.57 | 0.90 | 0.82 | 8.15 |
| Mean | 3.40 | 1.81 | 0.79 | 0.62 | 15.09 | 2.96 | 1.68 | 0.78 | 0.63 | 14.88 |

\* Animals are identified by the type of breed: Charolais (Ch) or Luing (Lu), and the type of basal diet: mixed (M) or concentrate (C).
\*\* CCC: Lin's concordance correlation coefficient.
\*\*\* CV<sub>RMSE</sub>: coefficient of variation of the root mean squared error.

Figure 5 shows the boxplots for the model parameters by treatment and basal diet. Table 3 shows the results of unpaired *t* tests at the 5% level testing for a difference in parameter means in concentrate vs mixed basal diets and rapeseed meal vs rapeseed cake. The model parameter *c* is significantly lower for a concentrate diet compared to a mixed basal diet, but parameter *d* is not significantly different (at the 5% level) for each diet. This is to be expected as *c* includes the methane yield factor that converts *u(t)* (DMI or IT) to *y*_*a*_*(t)*, whereas *d* is simply a specific rate constant related to gas release. While the methane yield depends on the level of concentrate in the diet, the concentrate level might not have impact on the rate of gas release (exhalation+ eructation). The level of dietary oil (oil and control) did not have significant effect on any parameters.

**Figure 5.**
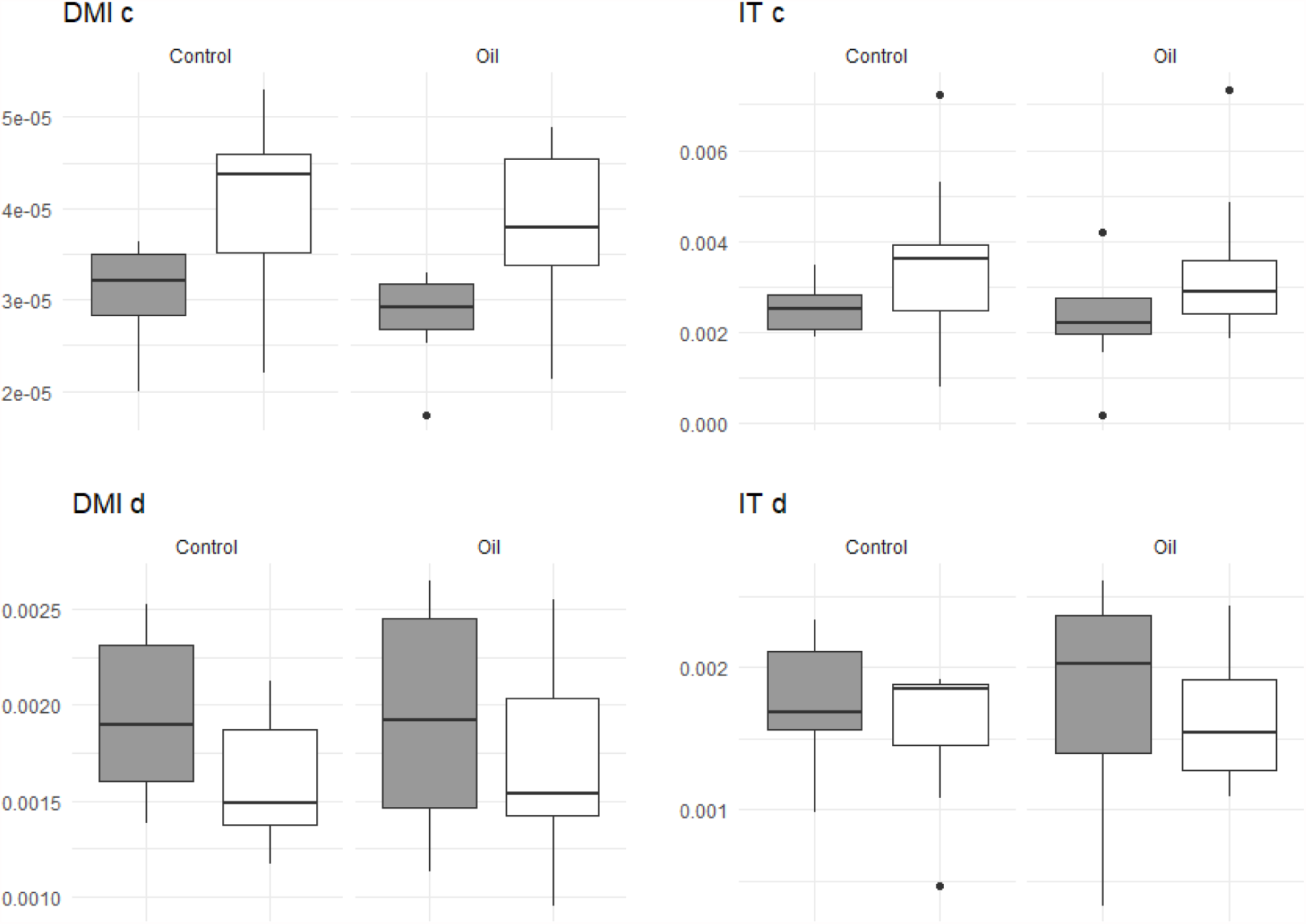
Estimated parameters of the model by treatment (control and oil). Grey boxes are for the concentrate basal diet and white boxes are for the mixed basal diet

**Table 3.**
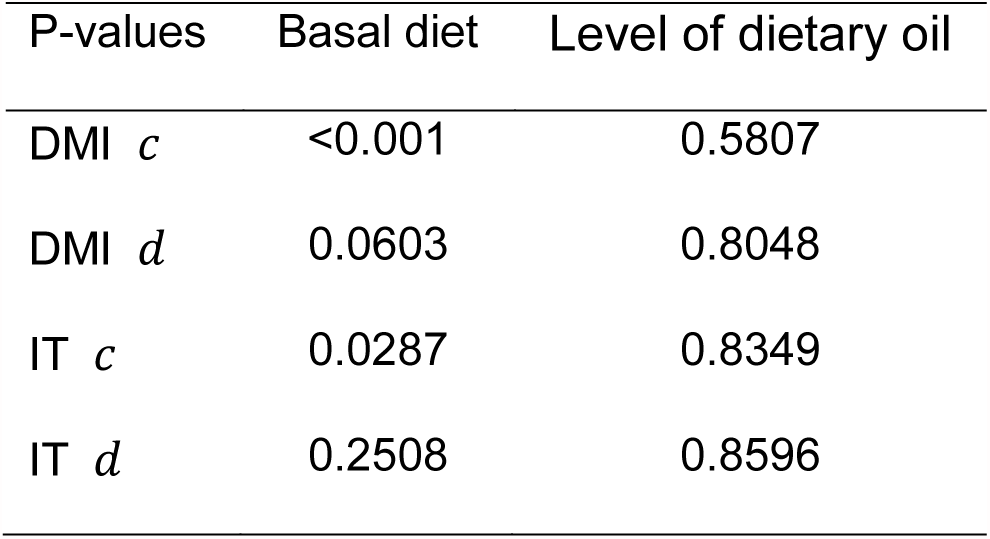
P-values when testing for a difference in parameter means between basal diet and level of dietary oil groups

Our software sensor provides satisfactory results for predicting the dynamics of methane production with similar levels of performance between DMI and IT as predictors. For DMI, the average CCC was of 0.79. For IT, the average CCC was 0.76. Interestingly, for 14 out of 37 dynamic data, the CCC for IT was higher than the CCC for DMI, indicating the great potential of using IT as predictor. When the software sensor was applied for predicting the daily average methane emission (Figure 4), the CCC was 0.99. On the basis of concordance analysis, our software sensor performs very well compared with reported literature results for methane proxies and predictive models (Wang et al., 2015, Negussie et al., 2017a, Niu et al., 2018).

## Discussion

The primary role of the feeding pattern on methane emissions in cattle (*Crompton et al., 2011*) motivated us to investigate the capability of predicting the dynamics of methane production from cattle using only time-series data of feeding behaviour (DMI or IT) *via* the construction of a software sensor. The objective of this construction was to develop a suitable tool for estimating methane that could be applied at large scale. The outcome of our work is encouraging to envisage a real-time implementation provided that accurate measurements or estimations of feeding behaviour are guaranteed, which is in compliance with other studies (Appuhamy *et al.*, 2016). Since the pattern of feeding behaviour is an individual trait among ruminants (Morita *et al.*, 1996, Giger-Reverdin *et al.*, 2012), individual characterisation of feeding patterns is central for producing individual estimations of methane by ruminants in a large scale context. Our study suggest that IT is a good predictor of methane emissions. The use of IT as predictor in our model relies on the assumption that the intake rate is constant across the day. This assumption applied to the data analysed here was shown to be adequate for methane prediction purposes, but additional data should be further analysed to assess the assumption robustness. For a real implementation at large scale, using IT as predictor instead of DMI has great advantages in terms of costs and setting. Successful results on real-time determination of IT by means of accelerometers (Oudshoorn *et al.*, 2013, Arcidiacono *et al.*, 2017) are encouraging to make of our software sensor a feasible and low cost solution for on farm applications in the future. Recently, it has been suggested that, for a stable management of feed allocation, the diurnal pattern of methane is constant over time (Bell *et al.*, 2018), which, with respect to our modelling work, translates into a constant diurnal feeding behaviour pattern. Accordingly, monitoring IT offers an opportunity not only to predict methane emissions but also a tool to characterise individual normal feeding patterns. By this, it will be possible to signal when an animal exhibits a different pattern from its normal pattern. This change of pattern could be associated to a perturbation, providing useful information for timely interventions.

Finally, a software sensor is meant to operate in real time using on-line measurements. In this work, however, our analysis was developed off-line. Further work needs to be carried out to evaluate the developed software sensor in real-time by integrating accelerometers for IT estimation. Running the software sensor requires prior calibration of the model parameters. This calibration can be performed, for example, by the GreenFeed system given its reliability for determining methane emissions on farm (Doreau *et al.*, 2018), provided an adequate setting (Renand and Maupetit, 2016). Once the model parameters are estimated, the software sensor can be applied by setting an initial condition for methane production. This initial condition has a strong impact on the amplitude of the methane emissions pattern. In our study, we extracted the initial condition from the experimental data. For real implementation, it would be recommended to start the software sensor at a moment when the methane production is close to the basal production of methane (*e.g.* before a meal). From the experiments analysed in this study, the basal production of methane was between 0.015 and 0.13 g/min. The strategy of starting the software sensor at a moment where the methane production is close to basal production reduces the impact of a wrong choice of initial condition. Since the methane emission pattern might change over age and physiological state of animals (*e.g.,* lactation stage for dairy cattle), adjustment of model parameters might be required when appropriate.

## Acknowledgements

We thank Dr. John A. Rooke for helpful explanations on the respiration data from the study conducted at SRUC. We thank Dr. Glenn Marion (BioSS) for his valuable comments for improving the clarity of the manuscript. John Fredy Ramírez Agudelo is supported by a PhD grant of Colciencias (Colombia). BioSS and SRUC are funded by the Scottish Government through the Strategic Research programme of the Scottish Government’s Rural and Environment Science and Analytical Services Division (RESAS). Helen Kettle was also funded under the Scottish Government’s Strategic Partnership in Animal Science Excellence.

## Supplementary material S1

### Influence of the turnover rate of a respiration chamber on methane output

In a respiration chamber, the rate of methane produced (g/min) by the animal (y_a_) and the rate of methane production measured in the chamber (_c_) are related by the following mass balance model

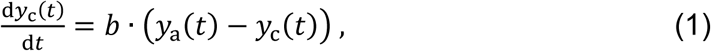

where *b* (min^-1^) is the turnover rate of the chamber. The magnitude of *b* determines how fast the trajectory of *y*_c_ will follow the trajectory of *y*_a_. The higher, the faster *y*_c_ converges to the trajectory of *y*_a_. As displayed in Figure S1, a wrong choice of the turnover rate will imply an important mismatch between the dynamics of methane produced by the animal and the dynamics of the methane flux of the chamber. An adequate turnover rate guarantees that the dynamic of *y*_a_ is mirrored by the dynamics of *y*_c_, that is that the approximation *y*_c_ ≈ *y*_a_ (used in this work) is consistent.

In theory, a very high turnover rate of the respiration chamber is ideal to capture the dynamics of methane produced by the animal. In practice, however, attention should be paid to very high turnover rates, since the gas flux at the outlet of the camber might be too fast for the gas analyser to produce consistent measurements. The rate of sampling of the gas analyser must be considered to select the optimal turnover rate. Since the overall efficiency of a respiration chamber depends on the extraction, conduction, and gas analysis (Gardiner *et al.*, 2015), both turnover rate and sampling rate are determining elements of the accuracy of respiration chambers for measuring methane emissions from livestock.

**Figure S1.**
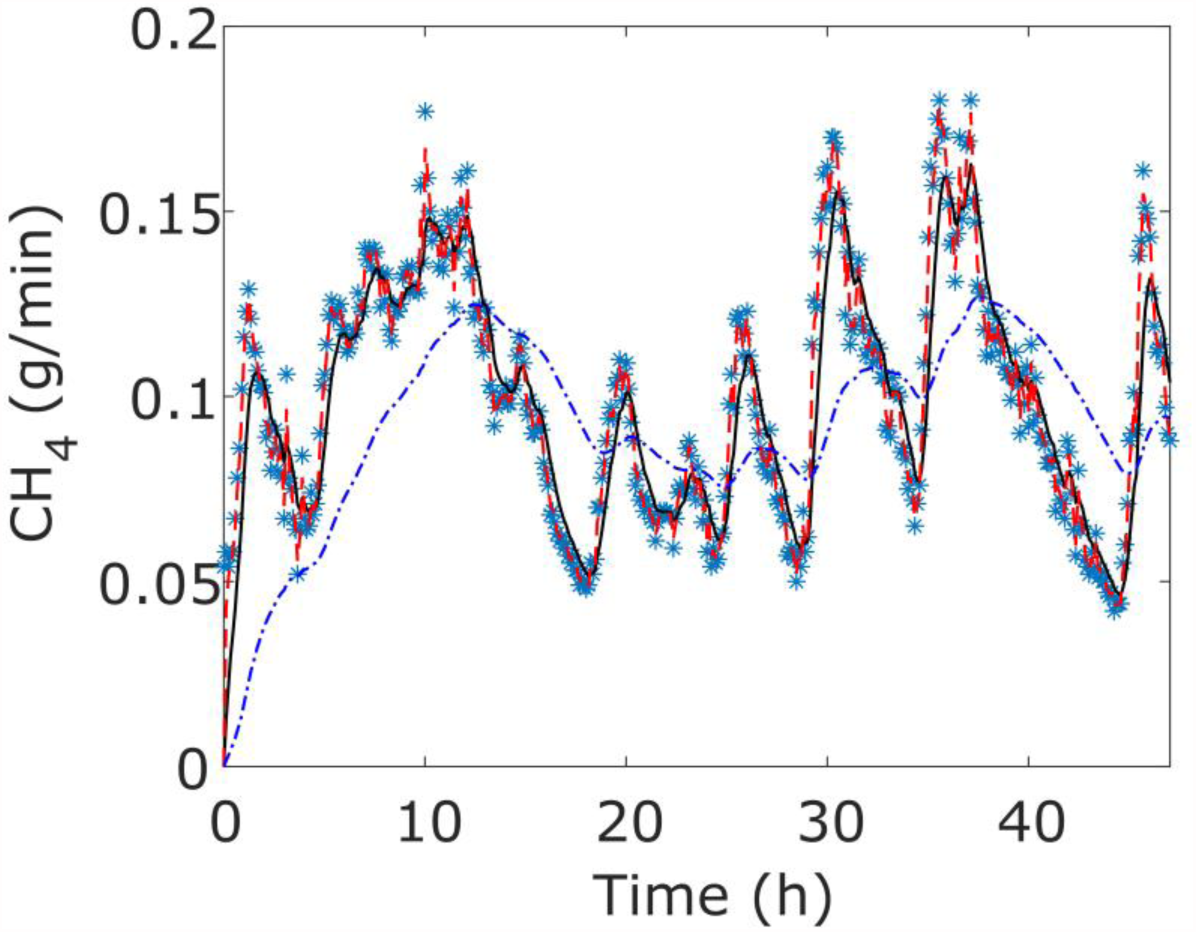
Simulation study. Virtual data of methane production by the animal (*) are compared with the output from a respiration chamber using Eq. (1) at three turnover rates: 0.004 min^-1^ (blue -.), 0.4 min^-1^ (red --), 0.04 min^-1^ (solid black line). The turnover rate of the respiration chambers used in this study was 0.04 min^-1^.

